# Body size but not age influences phototaxis in bumble bee (*Bombus terrestris*, L.) workers

**DOI:** 10.1101/673509

**Authors:** Michal Merling, Shmuel Eisenmann, Guy Bloch

## Abstract

We studied phototaxis, the directional movement relative to light, in the bumble bee *Bombus terrestris*. We first developed and validated a MATLAB based system enabling reliable high-resolution tracking of a bee and a measurement of her distance relative to a changing LED light source. Using this system we found in all our experiments that workers show positive phototaxis. The strength of the phototactic response was influenced by body size but not age, and this effect was significant when the light source was weak. In a separate experiment, foragers showed stronger phototactic response compared to nurses only in one of two trials in which they were larger and tested with weak light intensity. The evidence that phototaxis is associated with size-based division of labor in the bumble bee and with age-related division of labor in the honey bee, lends credence to response threshold models implicating the response to light in the organization of division of labor in cavity dwelling social insects.

## Introduction

Phototaxis is a behavior in which an animal moves towards (positive phototaxis) or away (negative phototaxis) from a source of elevated light intensity. The phototactic behavior is thought to be functionally adaptive because it regulates light exposure and facilitate orientation in space. For example, positive phototaxis may facilitate foraging behavior or escape in flying animals (e.g., Minot 1988; Reisenman et al.,1998), and negative phototaxis may protect animals by guiding them towards places hidden from predators (Hunte & Myers 2007; Minot 1988), or direct them towards burrowed nutrients (Sawin *et al.* 2004). Phototactic behavior is sensitive to light intensity and is commonly used to assess the animal capacity to discriminate between levels of light intensity. Pioneering studies with fruit flies showed already early in the 20^th^ century that strong light can inhibit the phototactic response that is induced by weaker light intensities (Carpenter 1905). Concerning the underlying mechanisms, phototaxis is one of the first behaviors for which mutations were identified (McEwen 1925). However, the phototactic behavior is not hardwired and is modulated by external factors such as temperature and odors (Shimizu and Kato 1978; Inoko et al. 1981), and internal factors such as diet, nutritional state, the endogenous circadian clock or mating state (Barker and Cohen, 1971; Inoko et al. 1981; Reisenman et al., 1998; Mazzoni *et al.* 2005; Bernadou and Heinze 2013). The phototactic behavior is also commonly modulated during development. For example, bird chicks switch from negative to positive phototaxis along the time they are ready to fledge (Minot 1988), and in Drosophila, young larvae show negative phototaxis which later changes to positive near the end of the third instar and at the final instar. During this period the larva prepares for pupation seeking dry surfaces, rather than the moist depths of fruits (Sawin *et al.* 2004; Mazzoni 2005).

The honey bee has been one of the first model systems with which to study phototactic behavior and its underlying mechanisms (Bertholf 1927). As in other animals, phototaxis in honey bees was shown to be sensitive to light intensity (Bertholf 1927; Kaiser et al., 1977; Menzel & Greggers, 1985). The phototactic behavior in bees can be mediated by multiple photoreceptors and is not restricted to a narrow wavelength (Kaiser et al., 1977; Menzel & Greggers, 1985). In addition to the compound eyes, there is also evidence suggesting that the ocelli are important for proper phototactic behavior in honey bees (Kastberger, 1990). The honey bee provides some of the most striking examples for developmental modulation of phototaxis. First, the developmentally-determined caste of female bees affects their phototactic behavior during the adult stage. Whereas phototaxis is negative in queens, it is strongly positive in foraging worker bees. Additional modulation occurs during the lifetime of worker bees. The phototactic behavior of the worker changes with age and experience (e.g., orientation flights; Vollbehr, 1971). Later studies further linked this association to age-related division of labor. Young workers typically tend brood inside the dark nest and show weaker phototaxis compared to their older sisters that typically perform foraging activities outside the nest (Ben-Shahar et al., 2003). Pharmacological studies support this association by showing that treatment with neurochemicals that modulate age-related division of labor or show increased amounts with age, such as cGMP, seretonin (5-HT) and tyramine, can enhance the positive phototactic behavior as expected for a trait that is associated with foraging activities (Ben-Shahar et al., 2003; Thamm et al., 2010, Scheiner et al., 2014). There is also evidence suggesting an even finer task-related modulation of the phototactic behavior, which is associated with the specialization of foragers in pollen or nectar collection (Scheiner et al., 2014, Erber et al., 2006, Tsuruda & Page, 2009). However, given that the differences between pollen and nectar foragers were not consistent across studies, this interesting hypothesis needs to be further tested.

The association of task performance and phototaxis in honey bees is thought to be functionally significant because strong positive phototaxis can guide bees towards the nest entrance and position them where they may be induced to forage by exposure to additional foraging related stimuli. Weak or negative phototaxis may be advantageous for nurses because the brood is typically located deep in the dark cavity of the nest (Ben Shahar et al. 2003). These observations for honey bees are consistent with response threshold models stating that variation in internal thresholds for responding to task-related stimuli contribute to the division of labor among workers in a colony (Beshers and Fewell, 2001; Beshers et al. 1999; Bonabeau and Theraulaz 1999). However, this hypothesis has not been tested in relation to phototaxis in other species of social insects. Here we study phototaxis in the bumble bee *Bombus terrestris*, in which the division of labor relates to body size, with relatively little influence of age (Yerushalmi et al. 2006). Body size in bumble bees is also correlated with anatomical, physiological and morphological differences that appear to improve foraging performance (reviewed in Chole et al. 2019). For example, large workers have larger eyes (Kapustjanskij et al. 2007; Spaethe and Chittka 2003), and a greater density of antennal sensilla (Spaethe et al. 2007), which are associated with better visual and olfactory accuracy. Their brain is larger (Mares et al. 2005; Riveros and Gronenberg 2010) and they perform better in some learning assays (Riveros and Gronenberg, 2012; Worden et al. 2005). They also have more cells expressing the circadian neuropeptide Pigment Dispersing Factor (PDF; Weiss et al. 2009) and stronger and earlier circadian rhythms (Yerushalmi et al. 2006). It has been also suggested that smaller bees are better suited to performing some in-nest activities (Brian 1952; Couvillon and Dornhaus 2010). This individual variation is largely independent of genetic effects because workers in a bumble bee colony are typically the daughters of one singly mated queen and are therefore closely related. Given this association between body size, function and task performance, we hypothesized that large bumble bee workers will have stronger phototaxis compared to their smaller sister worker bees. Age is also an important factor that commonly influence phototactic behavior (see above), and in bumble bees, the propensity to forage increases with worker age (e.g., Yerushalmi et al. 2006). However, given that age has less influence on task-performance, we hypothesized that it will have a weaker effect, if at all. To test these hypotheses we developed a new phototaxis monitoring system, allowing continuous tracking of the phototactic response of individually isolated bumble bees exposed to various light intensities. We next used this system to compare the phototactic behavior of nurse and forager worker bees differing in body size or age. Our results suggest that body size but not age is positively associated with phototaxis.

## Materials and methods

### Bumble bees

We purchased *Bombus terrestris* colonies from Polyam Industries, Kibbutz Yad Mordechai (Israel). Colonies were obtained a few days up to two weeks after the emergence of the first worker and contained a queen, 1-50 workers (varied with the experiment, see below), and brood at all stages of development. We housed the colonies in wooden nest-boxes (30 × 23 × 20 cm) with a transparent Plexiglas cover. We performed all treatments and inspections under dim red light which bumble bees do not see well (Peitsch et al., 1992). We measured the length of each forewing twice as an index for body size (Yerushalmi et al. 2006).

### Monitoring phototactic behavior

The phototaxis monitoring system included a stand with a perpendicular wooden board to which we fixed a removable cage holder. The cage holders were made of 5 mm wide foamed cardboard (Kapa), and each contained four parallel slots into which we placed the monitoring cages (Fig. 1). The distance between the cages was 5 cm. The monitoring cages (30×5×5 cm) were made of a similar foamed cardboard and had front and sidewalls made of a transparent glass. In order to optimize contrast between the bee and the background, we used white foamed cardboard for the posterior part of the cage that faced the camera. The bottom and top walls were covered with opaque black cardboard to minimize exposure to light from the other cages. A white LED light with spectral wavelength ranging from 440 to 700nm (main peak at 460 nm, and a lower one at ∼550) was placed in each side of each of the four cages (**Fig. 1**). During the operation of the LEDs, the temperature varied by ±1° C. The LED lights were tested with a Polaroid polarizer to confirm that LEDs do not produced polarized light. We positioned the LED lights such that they faced a 40-degree angle downward. This positioning resulted in the light spot hitting a point next to the cage’s edge, with no bleaching to adjacent cages. To measure illumination intensity and spectrum we placed the sensor of an Ocean Optic Spectrometer inside the cage in a distance of 1.5 cm from the LED light. We also used Walz ULM-500 photometer to more precisely measure the light intensity inside the monitoring cages. A stand to which we fixed an infrared sensitive camcorder (Sony TRV 75E) was placed at a distance of 1 meter from the monitoring arena; the height of the camcorder was similar to that of the board with the cages (Fig. 1). The camera produced dim infrared light that illuminated the arena during the experiment. We conducted the phototaxis experiments inside a dark environmental chamber (28 ± 1°C; 50-60% relative humidity).

**Figure 1.**
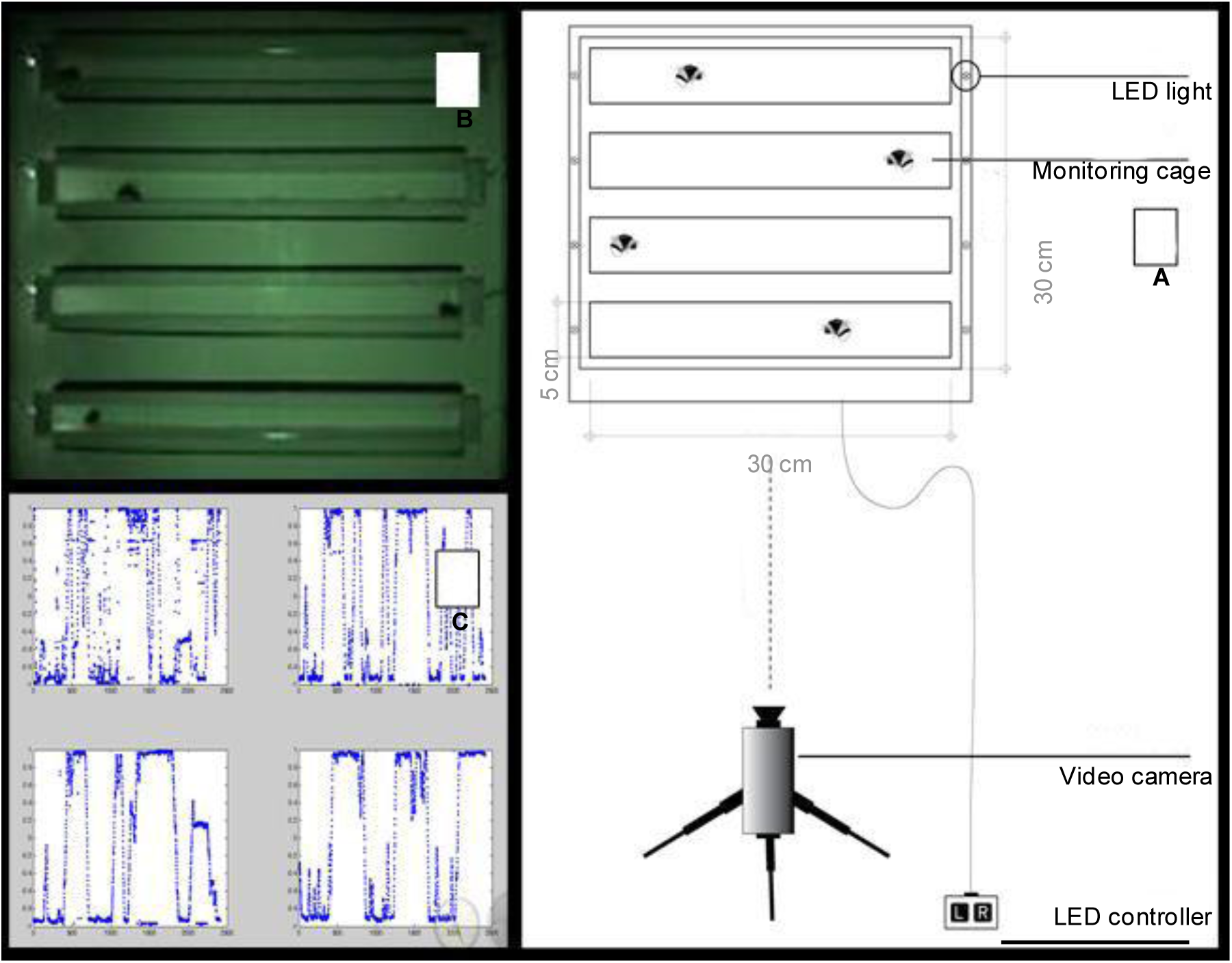
Setup for monitoring the phototactic response of insects. (A) A general scheme of the monitoring system. (B) A photo showing four cages, each with a single bee. The LED light on the left side of the cages is on. C) Representative traces produced by the tracking algorithm.

### Algorithm for automatic data acquisition and for phototaxis analysis

For the automatic monitoring of worker position relative to the LED light source, and for the analysis of the phototactic response, we developed a custom-made tracking code using MATLAB (version 7.0; the Math Works, Natick, Massachusetts, USA). The code identified an object with strong contrast relative to its background within a defined arena; each of the four cages was defined as an arena. The algorithm identifies the object (bee), tracks its location (center of mass), and records its position into a text file. It is also possible to divide an arena into distinct “zones” and to calculate how much time the object (bee) stayed within each zone (see *“Validation of the automatic phototaxis monitoring system” below)*.

We first converted the video records from the experiments into digital (AVI) formats with Pinaccle Studio 9 (Pinnacle, Systems, Mountain View, California). Next, we used the AVIedit software (AM Software, Russia) to convert the digital video file into a series of JPEG digital 720 × 576 pixels photos (12.5 frames/ second). In order to minimize the influence of background noise we used Adobe Photoshop 7.0 (Adobe Systems Inc, San Jose, California) to digitally remove the bees from a representative photo. This way we obtained a photo of the background. This background photo, which included the image of the cages but not the bees, was subtracted from all the digital photos (with bees) of the same experiment. We use the Image J image analysis software (US National Institutes of Health, Bethesda, Maryland, USA) to record the X, Y coordinates defining each one of the arenas (cages), and prepared two text files. In the first, we recorded for each frame the position (right or left) and state (on or off) of the light sources for each of the four arenas. The second included the following information: photograph resolution, range of frames in which the light is on, range of frame in which the light is off, X Y coordinates for the object (bee) in each arena, and the identity of the focal bee (i.e., forager or nurse, large or small, young of old).

The algorithm was composed of several scripts. In the main script *(mmain*) the software removes the background photo (without the bees, see above) from each frame in the file and performs an integral of the contrast on the Y-axis for each one of the arenas using a predefined coordinates. The second script *(getbee) extracted the* bee position on the x-axis for each frame as the average of the two most similar estimates out of three possible position estimates that were calculated. The first was simply the X value corresponding to the highest integral on the Y-axis. Given that the integral over the Y-axis sums the entire range of the cage, the shadow of the bee may be identify as the stronger contrast when the bee stands next to the light source. Therefore, the second and third estimates were designed to correct for this potential bias. The second estimate employs a specified script to identify the pixels with the sharpest change (the integral, calculated using the script “*deriv”*) in contrast value relative to neighboring pixels on the x-axis – this produced two points corresponding to the front and back parts of the bee. The position of the bee is calculated as the median between these two points (the “center” of the bee). The third estimate also uses the *“deriv”* script but calculates an accumulative integral. This estimate was specifically useful for cases of high background noise (for example, when the bee stands next to the light and causes apparent blinking in light intensity. An addition script (*DecideX)* compares the three position estimates and send to the main script the average for the two most similar estimates (see Supplementary Information for more details).

The positions of the bee at each frame were used to calculate the total path length she made during the experiment and the Phototaxis Index (PTI). The PTI was calculated by assigning for each frame one of three possible light position (“L” in Equation 1) values: -1 when the light is on the left side of the arena; 1 when the light is on the right side of the arena; 0 when the light is off on both sides. The position of the bee on the x-axis (“X” in Equation 1) was normalized such the -1 ≤ X ≤ 1, with -1, and 1 assigned to the left and right edges of the cage respectively. Equation 1 was used for calculating the PTI, in which n is the total number of frames analyzed by the algorithm for a specific bee:

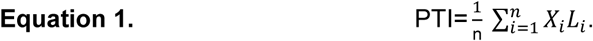

The PTI value varied between 1 and – 1, with positive values indicating positive phototaxis (i.e. the bee is close to the light), and negative values indicate negative phototaxis (i.e. the bee is far from the light). We omitted from the phototaxis analyses bees that were not active or show very little levels of activity during the experiment.

### Validation of the automatic phototaxis monitoring system

To confirm that our algorithm reliably capture the bee position we compared the automatically collected data with the actual path for ten bees monitored in different experiments (6 in high light and 4 in low light intensity). To determine the actual position of the bee, we used the ImageJ image analyses software to analyse the first 400 frames in each of these 10 records. We inspected every forth frame in the set of the first 400 frames and precisely determined the position of the bee center on the X-axis for each frame. We recorded the bee position (in pixels) for each of the 100 frames and normalized the X values such that -1 ≤ X ≤ 1, similar to the normalization of our monitoring algorithm. Using this information, we calculated the correlation coefficient between the automatically and manually estimated paths. We also used Equation 2 to calculate the percentage error for the automatically collected path.

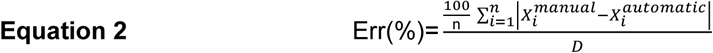

Where n is the number of frames, Xi^Manual^ is the normalized manually-determined position of the bee, Xi^automatic^ is the normalized software-determined position of the bee, and D is the length of the cage. D = 2 because after normalization the X values ranged between -1 ≤ X ≤ 1.

In a second set of validations, we compared the manual analyses of the phototactic response for the bees investigated in Experiment 1 in which the bee position was assigned to one of five zones (rather than a continuous measure as above). We first captured the video file into the computer (with Pinnacle Studio 9), and then divided each arena (cage) into five similar size zones ranging from the illuminated to the dark side of the cage. For each bee, we recorded the number of frames in which it was detected in each of the five frames. As an index for the phototactic response of the bee, we divided the number of times the bee was observed in the zone next to the light source by the number of frames she was in the zone in the opposite side of the cage. In this index, that we termed the “Light/ Dark Index” values >1 indicate positive phototaxis, and values <1 indicate negative phototaxis.

### Experiment 1. The influence of worker task on the phototactic response

Incipient bumble bee colonies were obtained a few days after the emergence of the first worker, and contained approximately 10 workers and brood at all stages of development. Immediately after receiving a colony, we marked all the workers with individual numbered tag (Graze, Germany), and transferred the entire colony into an observation hive made of transparent Plexiglas (32×24×13cm, described with more details in Yerushalmi et al. 2006). The observation hives have a cover with an opening allowing the removal and introduction of bees, and a cardboard bottom. We placed the observation hives on shock absorbers to minimize vibrations to which bumble bees are very sensitive. The observation hive was placed in an environmental chamber (28 ± 1° C; 50-60% relative humidity). All colony manipulations and observations were done under dim red light. During the first few days, the colony was fed *ad libitum* with commercial syrup (Polyam Pollination Services) and fresh pollen (collected by honey bees). The nest box was connected to the outside by a clear plastic tube (length = ∼1 m, diameter = 2 cm) allowing the bees to freely forage in the area surrounding the Bee Research Facility. Food provisioning was gradually reduced and eventually stopped when a focal colony contained 10 - 30 workers (7-13 days from the emergence of the first worker). However, in cases in which the colonies did not store sufficient nectar or pollen, we supplemented the nest as needed in order to prevent stress response (e.g., larval culling) by the nest bees. We performed detailed daily observations starting approximately a week after connecting the colony to the outside. Each day, we observed the entrance of the colony between 06:30 – 08:30 during the morning, and between 17:00 – 18:00 during the afternoon, for a total of three hours a day, and recorded the number tags of bees returning from pollen or nectar foraging trips. Additional observations on the activity inside the nest box were performed daily between 09:00-10:00 at the morning, and 16:00-17:00 at the afternoon. During these observations, we recorded the following behaviors: entering the hive with pollen or nectar, building or manipulating wax cells, opening brood cells, inspecting egg cells, and feeding larvae (see Yerushalmi et al. 2006 and Shpigler et al. 2013 for more details).

We carried out the phototaxis experiments when the colony contained at least 30 active foragers. We classified a worker bee as a fo age if she was eco ded pe fo mi g ≥20 fo agi g t ips, a d o, or very few, brood care activities during the three days preceding the phototaxis assay. We classified a bee as a nurse if during this period she was recorded inspecting or manipulating brood cells, or feeding larvae, and had never been observed flying outside the nest. All the phototaxis tests were done over three days, always between 09:00 – 12:00. We made the experiments and observations at the same time of day in order to prevent confounding influences of time of day or circadian clocks. We collected foragers at the hive entrance, and a similar number of nurses from inside the nest. Two nurses and two foragers were assigned randomly to each of the four monitoring cages. To prevent a bias due to cage position, we confirmed that nurses and foragers were not placed more often at a specific cage (position). The cages with the bees were placed in the monitoring arena and we allowed the bees to acclimate for 10 minutes before beginning the phototaxis assay.

We performed two trials with bees from colonies ET and HT. The protocols for these two trials differed in several parameters. In the first trial with colony ET, we used a light intensity that ranged between ∼15 µmol/m^2^/sec (measured at the end of the cage closest to the lit LED) to ∼0.2 µmol/m^2^/sec (measured in the middle of the cage). These trials included six sessions in which the light is on in one side of the cage. Each session lasted three minutes and was followed by five seconds of darkness for the bee the recover. At the end of the day, we determined the size of the bees and returned them to the colony and the cages were cleaned with 70% ethanol. In the second trial with colony HT, we adjusted the protocol based on lessons learned from the first trial. We shortened the sessions to two minutes each (since it takes a bee only about five seconds to walk the entire length of the cage) and limited the number of sessions to five (lowering the duration of the entire test to about 10 minutes, instead of more than 18 minutes in the first trial). We also lowered the light intensity to 0.2 µmol/m^2^/sec (measured at the end of the cage closest to the lit LED), 5.7×10^−4^ µmol/m^2^/sec (measured at the middle of the cage).

### Experiment 2. The influence of body size on the phototactic response

The colonies from which we collected bees for this experiment were obtained a few days after the emergence of the first worker and contained a queen, 1-10 workers, and brood at all stages of development. The bees were marked and transferred to wooden cages (30 × 23 × 20 cm) with a transparent Plexiglass top. The colonies were fed ad libitum and had no access to the outside. Every day, we collected all the newly emerged bees (easily identified by their lack of yellow pigmentation), recorded their size (front wing length), and marked each individually with a number tag. We designated bees as “large” or “small” if they were larger or smaller than the median of their colony, respectively. We attempted to collect bees on the edges of the size spectrum (i.e., the largest and smallest in the colony) but the size comparison is nevertheless relative and not in absolute terms. To minimize possible effects of age, we tested only bees 2-5 days of age, and repeated the experiment with bees from five different colonies.

For each phototaxis assay, we collected two large and two small bees that were assigned randomly to the monitoring cages. The phototaxis assay included two parts; each part was composed of six sessions of one min each, separated by five min of darkness. In the first part of the test, we used low light intensity (0.2 µmol/m^2^/sec), and in the second part, high light intensity (15 µmol/m^2^/sec). The two parts of the test were separated by 15 seconds of darkness.

### Experiment 3. The influence of worker age on the phototactic response

The experimental design and phototaxis test were overall similar to Experiment 2. In each trial of this experiment, we compared the phototactic response of two young bees (2-5 days of age) and two older bees (16 – 19 days of age). We repeated this experiment with bees from three different colonies: EA, HA, GS (that was also used in Experiment 2; each bee was tested only once).

### Statistical analyses

We used parametric statistics for both the PTI and activity level for which we confirmed that the values fit normal distribution. For each bee, we calculated an average PTI value and therefore the average for each treatment group was the mean of the averages that is normally distributed. In Exp. 1 in which we compared nurses and foragers, we used two-side unpaired t-tests, and in Experiments 2 and 3, we used three-way analysis of variance (ANOVA) with treatment (size, or age), colony, and cage position as factors. Given that in all the experiments the cage position did not have a significant effect, we report below two-way ANOVA tests with treatment and colony source as factors. For the statistical analyses of the Light / Dark position in the first experiment, we used the non-parametric Mann-Whitney test.

## Results

### Validation of the automatic phototaxis monitoring system

Our custom-made algorithm for automatically tracking the bee position produced a track that was very similar to that actually made by the bee (as determined by manual inspection of the bees position; Fig. 2). The percentage error (see Material and Methods for details) was very low under both low light (mean %Error = 0.71%, range = 0.45 – 0.84%), and high light (mean %Error = 1.81%, range = 1.13 – 2.27%; Fig. 2C). The manually and software determined tracks were highly correlated in all 10 tracks we analyzed, with the software produced record explaining in average 96% of the variation in the bee position (linear regression analyses; P< 0.001 for all analyses; R^2^ range = 0.809 – 0.998; Fig. 2).

**Figure 2.**
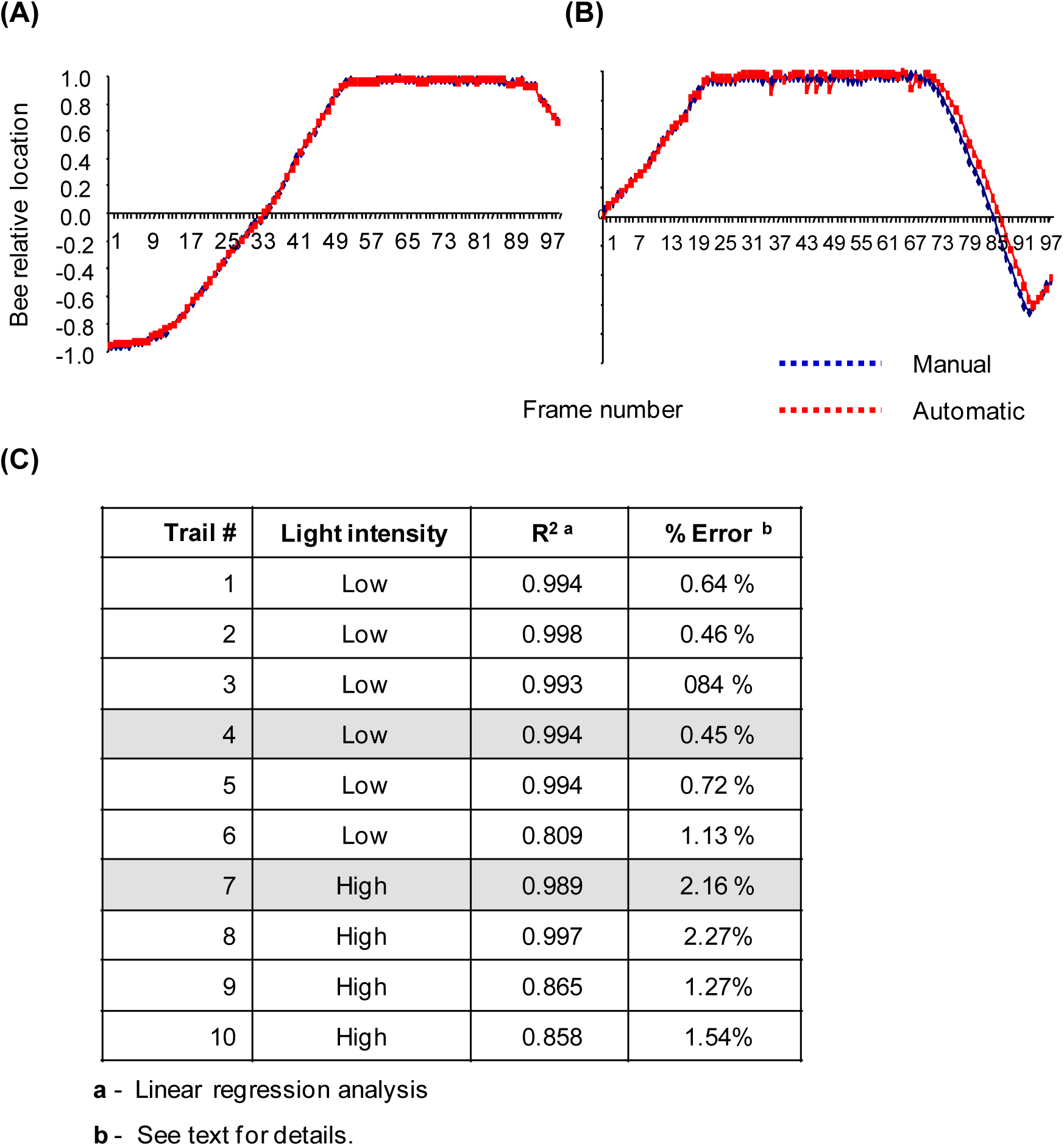
Validation of the automatic phototaxis monitoring system. We video-recorded ten bees during the phototaxis assay and determined their position with the automatic tracking system and manually by visually examining each frame. We determined the position of every 4^th^ frame in the set of the first 400 frames (n=100 frames). The Y-axis is the normalized position of the bee. **(A)** and **(B)** – comparison of the automatically and manually recorded positions of two representative bees. **(A)** A nurse bees assayed with low light intensity (Trial #4 in the table below). **(B)** A young bee assayed with high light intensity (Trial #7 in the table below). **(C)** A table summarizing all 10 trials. The trials highlighted with grey background correspond to the plots shown in (A) and (B) above.

In the second validation experiment, we used the Light/Dark index and compared records that were determined manually to those measured with the automatic monitoring system for bees from colony HT in Exp. 1. The results of the two analyses were overall very similar with a more positive phototaxis for foragers than the nurses (compare Colony HT in Fig. 3A and 3B). However, the differences in Light/Dark values were statistically significant only for the data collected automatically (Fig. 3A, B; Mann-Whitney test, two-tailed, P=0.155, P=0.024, for the manual and software collected data, respectively). The analysis with the software-recorded data was probably more powerful because sampling rate was 25 times higher. Complementary analyses using the Phototaxis Index (PTI) produced overall similar results for the two colonies (Fig. 3C). These two validation experiments indicate that our custom-made algorithm reliably tracks the bee position relative to the light source and produces a reliable and precise estimation of the phototactic behavior of the bee.

**Figure 3.**
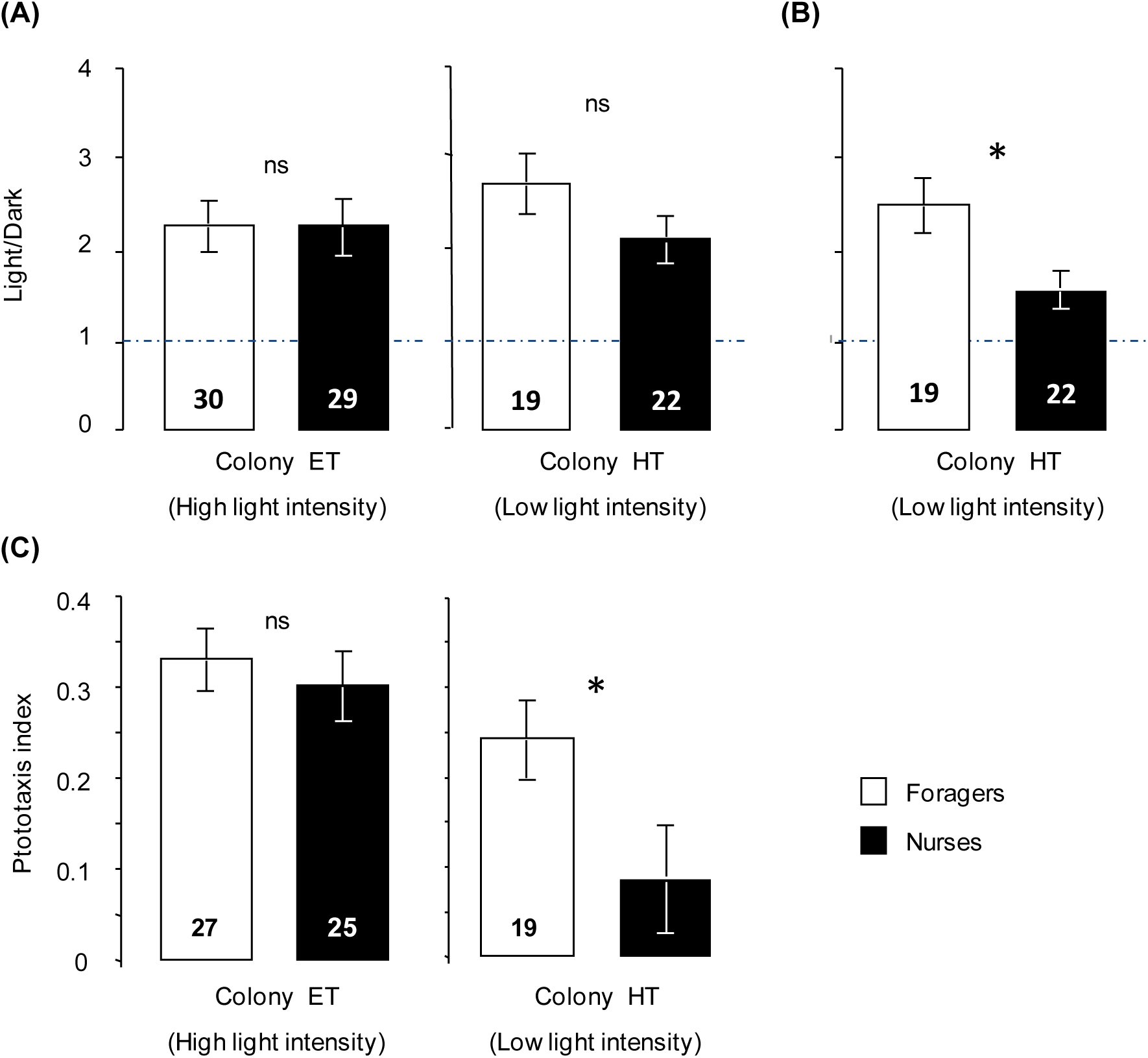
The phototactic response of foragers and nurses. **(A)** Manual determination of the bee position relative to the light source using the Light/ Dark index for the trails with colony ET (assayed with high light intensity) and colony HT (assayed with low light intensity). **(B)** Calculation of the Light/ Dark index based on the automatic phototaxis monitoring records for the trail with bees from Colony HT. **(C)** Determination of the phototactic response using the Phototaxis Index. The bars show mean ± SE, sample size within bars; * - P< 0.05. See text for more details and statistical analyses

### Experiment 1. The influence of worker task on the phototactic response

The two repetitions of this experiment differed in the details of the phototaxis assay protocol. In the first repetition with bees from colony ET, in which the bees were tested with strong light, both the nurses and foragers showed a positive phototaxis (single sample t-test compared with 0, P<0.001), that was similar for the two task groups (Fig. 3C; P=0.54; similar results were obtained using the manual Light/ Dark index, Fig. 3A). In this trial, the foragers and nurses did not differ in body size (data not shown) which is uncommon to the colonies that we obtained from commercial breeders. In the second repetition with bees from colony HT in which we used low light intensity, the foragers, but not the nurses showed a positive phototaxis (P<0.001, P=0.14, respectively; Fig. 3C). The PTI value was significantly higher for the foragers (unpaired t-test, P<0.05, see also Figs 3A and 3B for manual and automatic comparisons using the Light/Dark index, respectively). In this trial, the foragers were significantly larger than the nurses (Two-tailed t-test, P<0.001). In both repetitions the foragers and nurses did not differ in the level of locomotor activity (average speed; data not shown; unpaired t-test, P>0.5). This experiment shows that the phototactic behavior of bees can be modulated and suggests that body size is important because significant differences were obtained only in the trial in which foragers were significantly larger than nurses. To further asses the influence of body size on the phototactic behavior we next compared the phototactic response of same age small and large bees in colonies with no access to the outside.

### Experiment 2. The influence of body size on the phototactic response of workers

We repeated this experiment with bees from five colonies that were tested with both low and high light intensities. Both the large and the small bees had a positive phototaxis in the two light intensities ≤0.001 fo all the a alyses. Nevertheless, the positive phototactic response of large bees was significantly stronger compared with the small bees when tested with low light intensity (Fig. 4; two-way ANOVA with colony and size as factors; colony, P=0.41; size, P=0.003; interaction, P=0.14). A similar trend was seen when the bees were assayed with strong light intensity, but the influence of body size was not statistically significant (Fig. 4; colony, P=0.11; size, P=0.6; interaction, P=0.2). In two of the five tested colonies, the large bees showed higher levels of locomotor activity when tested with both low and high light intensity (data not shown; unpaired two-tailed t-test, P<0.05).

**Figure 4.**
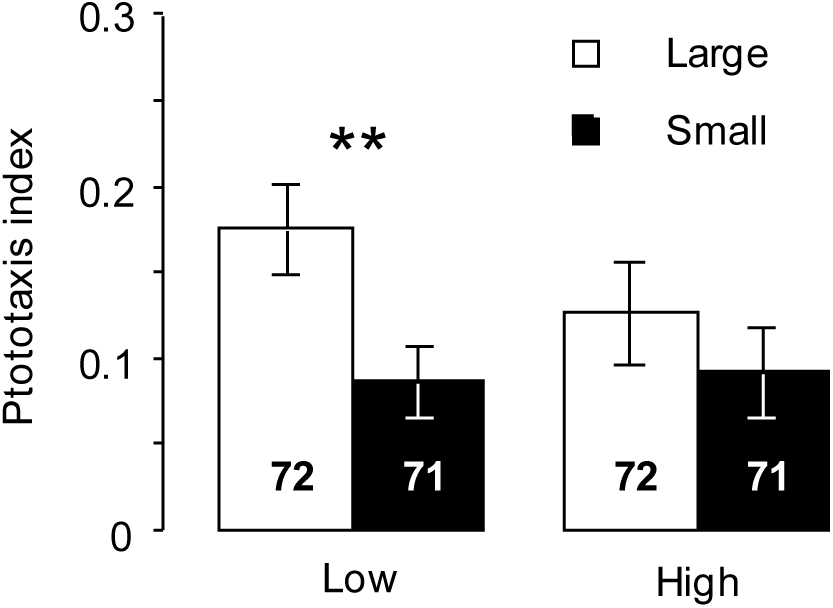
The phototactic response of large and small bumble bee workers. The plots summarize the results of five trials, each with bees from a different colony, that were assayed with low (left) and high (lright) light intensities. We used two-way ANOVA with Trial (colony) and body size as factors. ** - P<0.01. The Trial effect was not statistically significant in the assays with both low and high light intensity (P>0.1, see text for details). Other details as in Fig. 2.

### Experiment 3. The influence of worker age on the phototactic response

We repeated this experiment with bees from three different colonies that were assayed with both low and high light intensities. Both the young and the old bees had a positive phototaxis when tested in the two light intensities (P<0.01). Overall, there was no significant influence of age on the phototactic response of workers (Fig. 5.; Two-way ANOVA; age effect: P=0.49, colony effect: P=0.09, interaction: P=0.13). Young and old bees did not differ in their level of locomotor activity (Two-way ANOVA; low light intensity: P=0.97; high light intensity: P=0.61).

**Figure 5.**
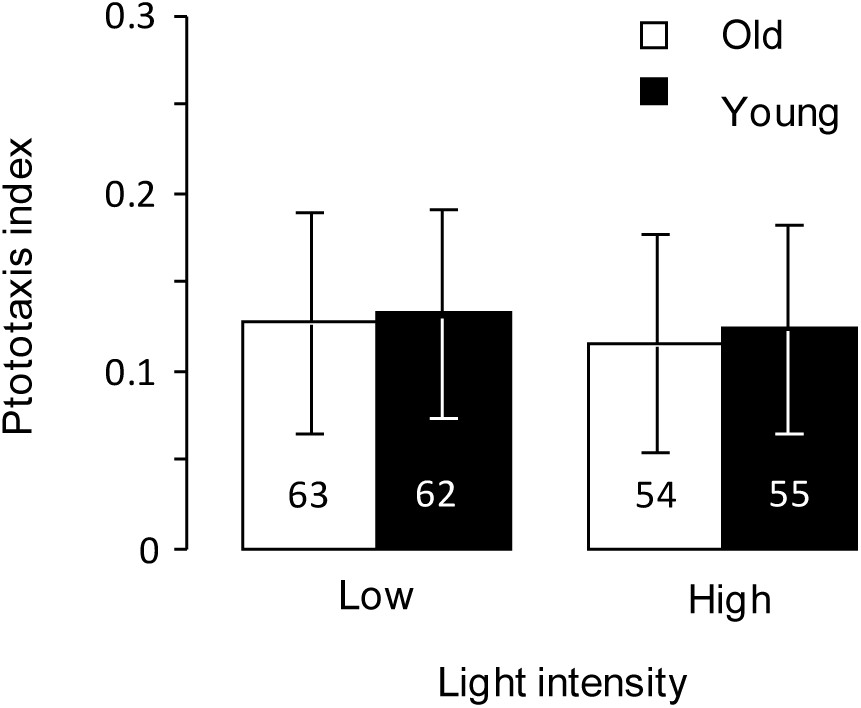
The phototactic response of young and old bumble bee workers. The plots summarize the results of three trials, each with bees from a different colonies, that were assayed with low (left) and high (right) light intensities. We used two-way ANOVA with trial (colony) and age as factors. The Trial effect was not statistically significant in the assays with both low and high light intensity (P>0.1, see text for details). Other details as in Fig. 3.

## Discussion

We developed and validated a MATLAB based tracking system enabling reliable high-resolution tracking of individually isolated bumble bees. To estimate the phototactic response we exposed the bees to LED lights that can emit light over a range of intensities. Our algorithm continuously tracks the bee location and calculates the distance between the bee and the light source, and thus provides precise quantification of the bee location relative to the light source. Importantly, we switched the light source between the two sides of the monitoring cage several times in each assay. This procedure enables us to separate the phototaxis response from simple directional movement, overall levels of locomotor activity, or arousal state. Our validation experiments in which we compared our tracking system to a manual frame-by-frame inspection of the bee location confirmed the precision and reliability of our system. This system can be adapted to study phototaxis in many other insect species.

Following the validation of our phototaxis analysis system, we used it to begin investigating the modulation of the phototactic behavior of bumble bees. We focused on three factors that were shown to affect phototaxis in animals; age, body size, and task performance (see Introduction for references). In all the trials in all three experiments, the bees moved toward the light, showing a positive phototaxis. However, the intensity of the phototactic response was modulated, with the stronger effect for body size. Larger bees showed overall stronger phototactic response compared with their smaller sisters. This difference was statistically significant only for bees tested with the lower light intensity. The effect of light intensity is not surprising, because the phototactic response of animals is typically influenced by the intensity of the light source (Carpenter 1905; Yamaguchi et al. 2011; Menzel and Greggers, 1985; Erber et al. 2006). Thus, although small bees can show relatively strong positive phototaxis when light intensity is high, their weaker phototaxis relative to that of large bee suggest that they have a higher response threshold. A better description of the effect of body size on phototaxis can be achieved by studies using a broader range of light intensities. By contrast, to the overall effect of body size, we did not find an affect of age. Phototaxis is commonly influenced by age, specifically in animals that show ontogenic changes in behavior (Minot 1988; Sawin *et al.* 2004; Mazzoni 2005). These include the honey bee in which older bees show stronger phototactic response (Vollbehr, 1971; Ben Shahar et al., 2003). In honey bee colonies, workers show a form of behavioral development that underlies colony level division of labor. Young bees perform mostly tasks inside the hive such as care for the brood, whereas older bees typically perform activities outside the nest such as guarding the hive entrance and foraging for rewarding flowers. It is assumed that a strong positive phototaxis helps guide foragers to the hive entrance and orient them outside, whereas weak or negative phototaxis is beneficial for nest bees that perform activities in the dark cavity of the nest (Ben-Shahar et al., 2003). Given that in bumble bees worker task relates to its body size, with only little effect to age (Michener et al. 1974; Alfrod 1975; Yerushalmi et al. 2006; reviewed in Chole et al. 2019), the association of strong positive phototaxis with body size in bumble bees and with age in honey bees is consistent with this hypothesis. An additional support for an association between task and phototaxis came from Exp. 1 in which we explicitly compared bees that specialized in nursing and foraging activities. Foragers showed stronger phototactic response only in the trial with bees from colony HT in which phototaxis was tested with the lower light intensity and in which the tested foragers were significantly larger than nurses. We do not know whether the lack of influence of task performance in the trial with colony ET is because this colony was tested with a stronger light intensity, because in this colony nurses and foragers did not differ in size (given that we seen many bee-eaters around our Bee Research Facility during this repetition, perhaps this colony suffers from higher predation pressure that forced also relatively small bees to forage), or because genetic differences between colonies ET and HT affect their phototactic behavior. Nevertheless, the results of the trial with colony HT strengthen the positive association between body size and phototaxis that we found in the second experiment. Additional studies with bees representing a diverse genetic background, that are beyond the scope of the current paper, should test whether and how foraging experience itself affects the phototactic response.

In a broader view, our findings are consistent with predictions of response threshold models for the division of labor in insect societies. Workers in social insect colonies specialize in different tasks such as foraging, brood care, guarding, or nest cleaning. This specialization forms a division of labor, which is one of the most important organization principles of insect societies (Wilson 1971). But how does a worker knows which task it should take? Response threshold models state that workers have internal thresholds for responding to task-related stimuli and that variation in thresholds to different tasks among workers in a colony generates a division of labor (Beshers and Fewell, 2001; Beshers et al. 1999; Bonabeau and Theraulaz 1999). For social insects such as the Western honey bee, bumble bees, and many species of ants that nest in dark cavities but forage under sunlight, the entrance light stimuli are associated with foraging outside the nest. Thus, response threshold models predict that individuals with strong positive phototaxis (i.e., lower response threshold to light) are more likely to approach the nest entrance and take foraging tasks compared to individuals showing a negative or weak positive phototaxis (Ben-Shahar et al. 2003; Erber et al. 2006). Given that in bumble bees, larger workers are more likely to take foraging tasks, response threshold models predict that they will be more sensitive to foraging related stimuli compared to their smaller sisters, even when they have no previous foraging experience. The finding of Exp. 2 showing that large bees that never forage outside the nest nevertheless showed overall a stronger positive phototaxis than smaller bees is consistent with this prediction. The influence of body size on phototaxis in the current study also adds to previous studies showing that large bees typically perform better in behaviors that are associated with foraging activities (Kapustjanskij et al. 2007; Spaethe and Chittka 2003; Riveros and Gronenberg, 2012; Worden et al. 2005; Yerushalmi et al. 2006; reviewed in Chole et al. 2019). Taken together, these studies suggest that developmental processes that affect that final size of the bumble bee larva (Shpigler et al. 2013; Chole et al. 2019) determine not only the size of the emerging bee, but also behavioral and physiological qualities that affect her propensity to specialize in a certain task.

## Acknowledgements

We thank Moshe (Muki) Nagari and Nir Keren for help with the light intensity measurements. This research was supported by grants from the US-Israel Binational Science Foundation (BSF, Contract Numbers 2003151, and 201788, to G.B.) and the US–Israel Binational Agricultural Research and Development Fund (BARD, project No. IS-5077-18 R, to G.B.).

## Authors Contribution

MM and GB conceived this research and designed experiments; MM performed experiments and analysis; SE wrote and validated the code for bee tracking and phototaxis analyses. MM and GB wrote the paper and participated in the revisions of it. All authors read and approved the final manuscript.

